# LucaCell: a sequence-centric foundation model for cross-species single-cell analysis

**DOI:** 10.64898/2026.09.08.750024

**Authors:** Yan Sun, Yong He, Minsi Ren, Yanhui Wang, Penghao Xu, Yong Hou, Yu Kang, Tingjun Hou, Jieping Ye, Huanming Yang, Zheng Wang

## Abstract

Single-cell foundation models have transformed transcriptomic analysis, yet most rely on fixed gene identifiers that limit transfer across species and data types. Here we present LucaCell, a sequence-centric foundation model that represents genes through pre-trained mRNA sequence embeddings rather than static gene annotations. Gene expression is discretized into bins and modeled with a Transformer encoder, enabling sequence-informed cell representation without a fixed gene-ID vocabulary. Pre-training on 85 million human and mouse single cells, LucaCell is evaluated on human, mouse and lemur gene expression profiles, human chromatin accessibility data, unaligned reads from more than 50 prokaryotic taxa, and five influenza A virus genomes. LucaCell enables manual-mapping-free cross-species cell type annotation and an alignment-free microbial embedding framework that simultaneously distinguishes bacterial species identity and intra-species physiological states. It also improves gene expression reconstruction by incorporating donor-specific exonic SNP information into mRNA sequence embeddings, and predicts cellular viral load across influenza A virus strains while highlighting infection-like transcriptional states in mock-infected cells. These results show that sequence-informed gene representation can improve the generalization of single-cell foundation models across species, data types, and predictive tasks.

## Introduction

Single-cell RNA sequencing (scRNA-seq) has transformed our ability to resolve the cellular composition and organization of complex biological systems. The rapid adoption of scRNA-seq has generated an expanding repository of data that now encompasses hundreds of millions of cells. Major international consortia and data initiatives, such as the Human Cell Atlas ^1^, the Tabula Sapiens ^2^, and the CZ CELLxGENE data portal ^3^, have consolidated this growing body of information and enabled systematic exploration of cellular states and regulatory programs. Deciphering gene expression networks within cells is essential for understanding fundamental biological processes, including cellular differentiation, organismal development, disease mechanisms, therapeutic responses, cellular reprogramming, and aging. Despite this potential, the high dimensionality, sparsity, and measurement noise of single-cell data, together with heterogeneity across samples, technologies, species, and data types, pose substantial challenges for traditional analytical frameworks. Current computational methods ^4–6^ still struggle to achieve unified integration of such large, multi-batch datasets, as they often rely on fixed mapping relationships that limit the transfer of information across batches, data types, and species.

To address these challenges, the field has increasingly turned to foundation models, particularly large-scale neural networks pre-trained on large transcriptomic corpora. By leveraging Transformer-based architectures, foundation models such as Geneformer ^7^, scFoundation ^8^, scGPT ^9^, and CellFM ^10^ can capture contextual gene-gene relationships and learn latent cellular representations that generalize across experiments. Despite this progress, most existing foundation models treat genes as discrete symbols or identifiers, which limits their scalability to novel species, unannotated genes, and non-canonical genomic regions, thereby restricting their broader applicability in biological prediction. Moreover, while some recent cross-species models such as TranscriptFormer ^11^ and Universal Cell Embeddings (UCE) ^12^ have incorporated protein-sequence embeddings, they do not capture untranslated regions (UTRs) that contribute to transcript-level regulation, and their reliance on amino acid sequences limit flexibility for modeling non-protein-coding contexts.

To address these limitations, we introduce LucaCell, a nucleic acid sequence-centric foundation model for single-cell analysis. Departing from traditional identifier-based approaches, LucaCell represents genes through pre-trained mRNA sequence embeddings rather than relying solely on static gene annotations. This approach maps each gene into a high-dimensional latent space based on sequence information rather than discrete labels, enabling sequence-based representation across species and data types without requiring manual gene mapping. Unlike discrete identifiers that treat each gene as an independent symbol, sequence embeddings preserve information about sequence similarity and functional relatedness, allowing the model to recognize homologous genes even when names or annotations differ across species. By integrating these sequence-level biological priors into a Transformer-based architecture, LucaCell learns representations that reflect sequence-associated information and support scalable single-cell analysis across diverse settings.

In this work, we show that LucaCell can: (a) support sequence-centric representation learning that reduces reliance on species-specific gene annotations, enabling cross-species comparisons without manual ortholog mapping; (b) learn a scalable representation of cellular heterogeneity through pre-training on 85 million human and mouse cells spanning 62 tissue origins and 18 disease conditions; (c) incorporate nucleotide-level sequence information that capture variation not reflected in protein-level aggregation; and (d) support virtual cell applications, including in silico perturbation modeling and cell-pathogen interaction prediction. Together, our results suggest that transcript sequence information can improve the transferability of single-cell foundation models across species and biological contexts.

## Results

### Overview of LucaCell

Gene representation plays a central role in single-cell foundation models. We developed LucaCell, a sequence-centric foundation model that represents genes through their intrinsic mRNA sequences, thereby reducing dependence on predefined gene annotations (Fig. 1a). To operationalize this design, each gene is encoded by an mRNA sequence embedding derived from the LucaOne foundation model ^13^, and expression values are discretized into bins for model input. This sequence-centric tokenization supports integration across species and data types without requiring a fixed gene-ID vocabulary, thereby enabling LucaCell to exhibit capabilities distinct from those of existing single-cell foundation models (Table S1). Using a Transformer-based encoder architecture, LucaCell learns contextual gene relationships through a masked gene modeling objective that predicts masked expression bins from local cellular context. To capture global cellular heterogeneity, LucaCell was successfully pre-trained on 85 million single cells, including 67 million human and 18 million mouse scRNA-seq profiles spanning 62 tissue origins and 18 pathological states (Fig. 1b, S1). This sequence-based framework enables LucaCell to be applied to traditionally challenging downstream tasks, such as cross-species cell type annotation, alignment-free cell embedding, variant-associated gene expression modeling, and cell-virus interaction prediction (Fig. 1c).

**Figure 1.**
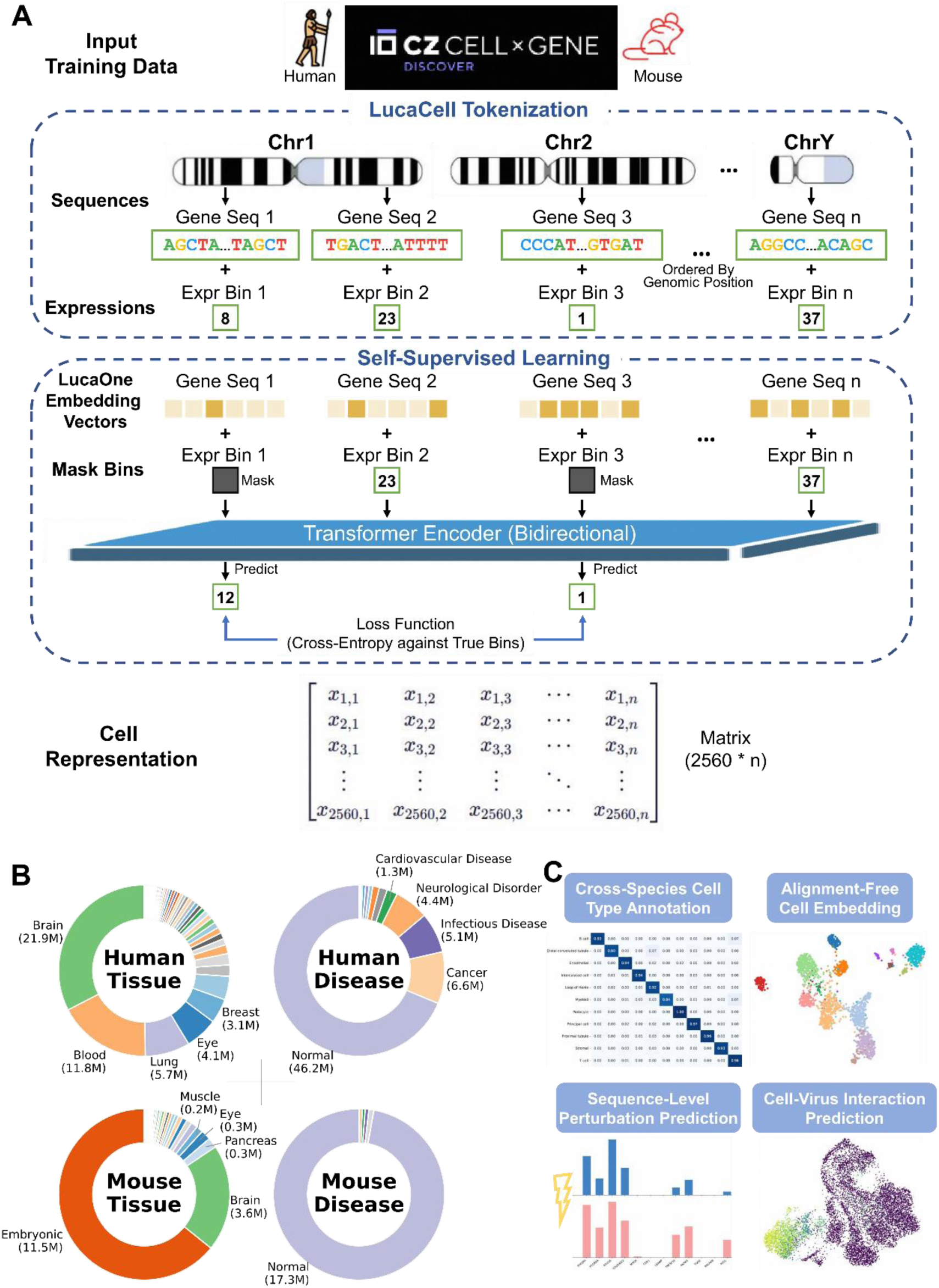
Sequence-centric tokenization and self-supervised pre-training of LucaCell. a, Schematic of the LucaCell tokenization and pre-training framework. Genes are represented by mRNA sequence embeddings derived from LucaOne and discretized expression bins. LucaCell is trained with a masked gene modeling objective to predict masked expression bins from cellular context using a Transformer encoder. b, Tissue and clinical composition of the 85-million-cell pre-training corpus. c, Sequence-informed downstream applications of LucaCell, including cross-species cell type annotation, alignment-free cell embedding, sequence-level perturbation prediction and host-virus interaction modeling.

### Cross-species and cross-modal cell type prediction

Accurately identifying cell types across evolutionarily divergent species remains a fundamental challenge, requiring computational frameworks to extract conserved gene regulatory programs from species-specific variation. To evaluate the cross-species and cross-modal generalization of LucaCell, we designed a transfer learning framework across four kidney datasets annotated with 11 shared cell types: human scRNA-seq and scATAC-seq ^14^, gray mouse lemur (Microcebus murinus) scRNA-seq ^15^, and mouse scRNA-seq ^16^, with the mouse scRNA-seq held out as a completely unseen test set (Fig. 2a). LucaCell yielded strong performance on seen species, achieving a test accuracy (ACC) of 0.937 (macro F1 = 0.921) for human scRNA-seq and 0.961 (macro F1 = 0.908) for lemur scRNA-seq. For the more challenging chromatin accessibility modality (human scATAC-seq), the model achieved a test accuracy of 0.653 (macro F1 = 0.516). On the completely unseen mouse dataset, LucaCell maintained moderate generalization performance (accuracy = 0.650, macro F1 = 0.493), indicating that sequence-informed representations support transfer across species.

**Figure 2.**
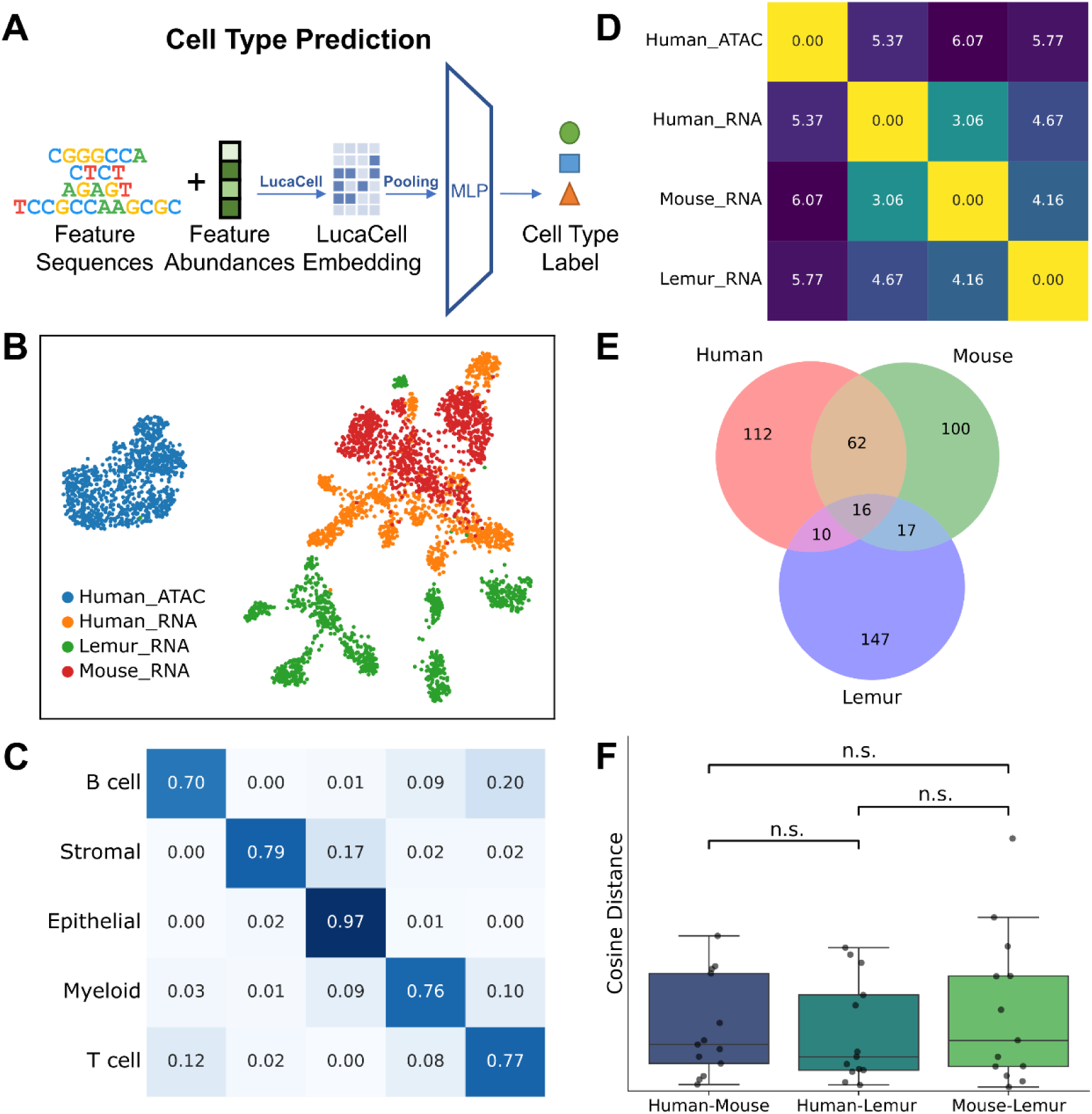
Cross-species and cross-modal cell type prediction and conservation analysis. a, Schematic of the sequence-aware transfer learning framework. Sequence-centric cell representations are processed through an MLP classifier to predict cell type identities across species and modalities. b, UMAP projection of LucaCell-derived cell embeddings, colored by species and modality. c, Confusion matrix showing cross-modal cell type prediction across five major cell types in human scATAC-seq data. d, Heatmap of mean pairwise Euclidean distances between cell embeddings across the species and modalities. e, Venn diagram showing the overlap of marker genes identified across all cell types in human, mouse and lemur. f, Pairwise distances between orthologous marker-gene sequence embeddings across species pairs. Statistical significance was assessed using a paired t-test; n.s., not significant (P > 0.05).

To investigate the structure of these representations across modalities and species, we projected cell embeddings from the pooling layer (the representation vectors) using UMAP. The projection showed that human scATAC-seq cells formed a distinct cluster separated from RNA-based modalities, while retaining cell type-specific structure (Fig. 2b, S2). When granular subtypes were aggregated into five major cell types, classification accuracy on human scATAC-seq data increased to 0.877 (macro F1 = 0.791; Fig. 2c), suggesting that LucaCell captures conserved cellular identity signals despite modality-specific noise. However, performance was lower for scATAC-seq subtype identification than for transcriptomic tasks, likely reflecting the sparsity and high dimensionality of chromatin accessibility data. Because the current implementation processed up to 8,192 ATAC peaks per cell without highly variable feature selection, subtype-level resolution remained limited.

UMAP visualization and Euclidean distance analysis showed that human and mouse scRNA-seq embeddings were more tightly integrated than human and lemur embeddings in this kidney-specific context (Fig. 2b, d), despite the closer phylogenetic relationship between humans and lemurs. To examine this pattern, we compared the transcriptional conservation and sequence similarity of orthologous genes across three species. Marker-gene analysis showed that human and mouse shared a larger set of orthologous markers across most cell types than either species did with lemur, indicating stronger conservation of renal transcriptional programs between human and mouse (Fig. 2e, S3). In contrast, pairwise comparisons of sequence embeddings for 13 highly conserved orthologous markers shared by all three species showed no significant differences in sequence-level distances among species pairs (Fig. 2f). Notably, when mouse gene sequences were randomly replaced with sequences from other mouse genes while keeping the downstream model architecture and expression bins unchanged, prediction performance dropped sharply (accuracy = 0.25, macro F1 = 0.05), indicating that intact sequence-derived embeddings are important for cross-species transfer. Together, these results indicate that sequence-derived gene representations provide the foundational basis for cross-species transfer, while the extent to which cells align to the same cell type across species is governed by the conservation of gene expression programs.

### Alignment-free embedding resolves microbial species and physiological states

Metagenomic studies have greatly expanded our understanding of microbial diversity, ecology, and function across complex environments. However, many existing approaches typically distinguish microorganisms at the species level, limiting their ability to resolve finer functional heterogeneity within microbial communities. Single-microbe RNA sequencing (smRNA-seq) offers the potential to capture functional and physiological heterogeneity at the level of individual microbes^17–19^, but conventional workflows remain heavily dependent on reference genomes and alignment-based pipelines. This dependence restricts their scalability, particularly for uncultivated or poorly characterized species. To address these limitations, we trained a downstream classifier initialized with LucaCell embeddings to predict microbial taxonomic and functional states directly from RNA sequencing reads without genome alignment (Fig. 3a). For taxonomic classification, we trained the downstream model on a 54-class task curated from a human gut smRNA-seq dataset ^20^, comprising the 53 most prevalent species and an ‘unidentifiable’ category. The model achieved a test accuracy of 84.1% (macro F1 score = 0.78). Unsupervised clustering of embeddings from held-out cells separated different microbial species into distinct clusters (Fig. 3b). In addition, some species, such as *Neobittarella massiliensis* and MGYG000004894, formed multiple subclusters, suggesting that the model may capture intra-species heterogeneity.

**Figure 3.**
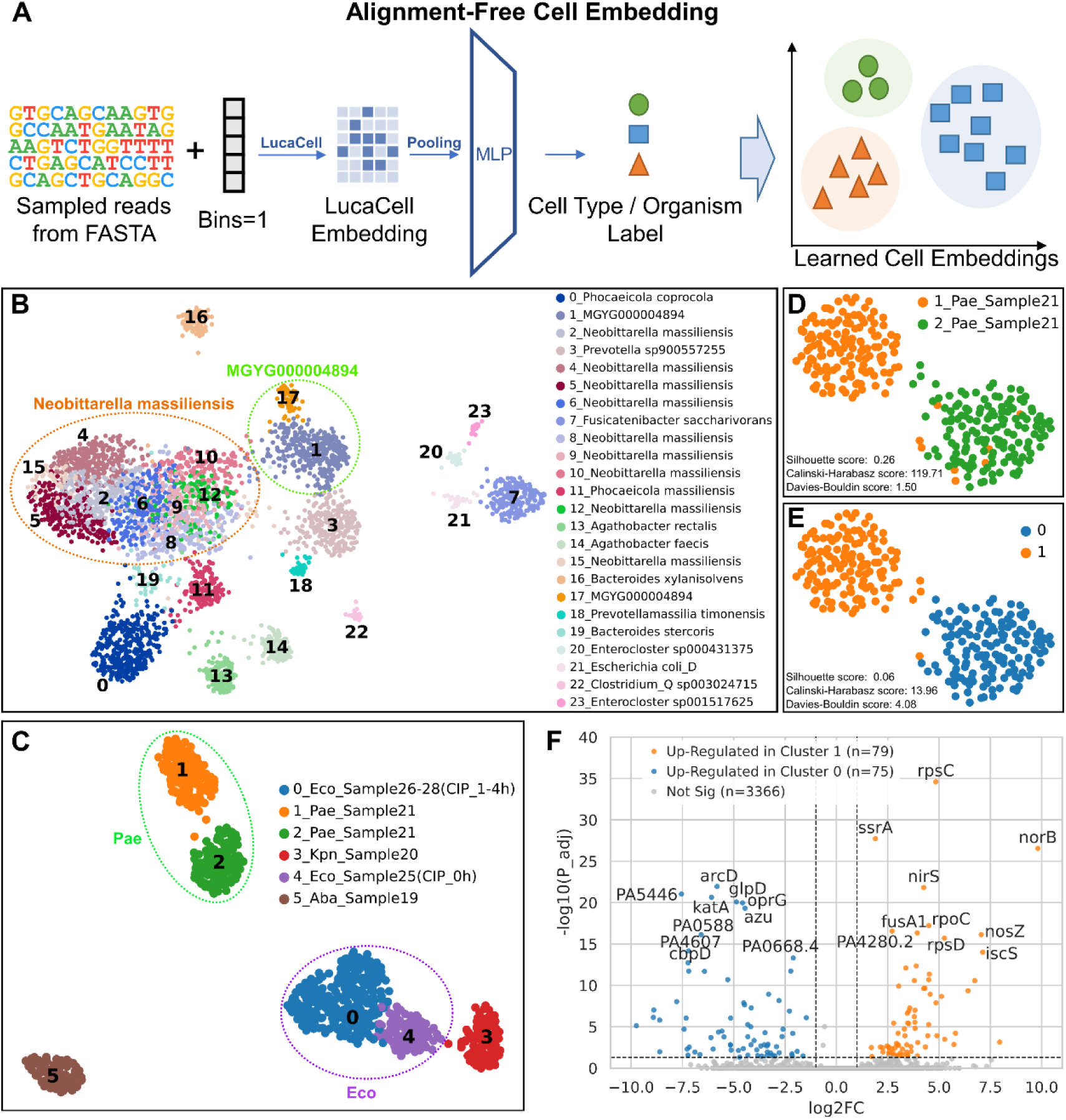
Alignment-free microbial representation resolves taxonomic identities and functional states. a, Schematic of the alignment-free microbial representation framework. Instead of aligned gene-count matrices, LucaCell processes raw sequencing reads directly. For each single-microbe, up to 8,192 reads are randomly subsampled, with their expression bins uniformly set to 1 to encode read presence (Methods). b, UMAP projection of LucaCell embeddings from human gut microbial smRNA-seq data. c, UMAP visualization of raw-read embeddings from an independent validation cohort containing unseen species, resolving taxonomic identities and drug-induced physiological transitions within Eco subpopulations. d, UMAP projection of the Pae gene expression matrix, colored by alignment-free clusters identified in c. e, The same UMAP projection as in d, colored by clusters derived from gene expression matrix, showing agreement between read-based and expression-based embeddings. f, Volcano plot of DEGs between Pae clusters 0 (blue) and 1 (orange), as shown in e. The top 20 DEGs with the lowest adjusted P-values are labeled.

To examine whether these raw-read embeddings reflect physiologically relevant transitions, we applied the trained model to an independent bacterial culture smRNA-seq dataset ^21^ comprising four species: *Acinetobacter baumannii* (Aba)*, Escherichia coli* (Eco)*, Klebsiella pneumoniae* (Kpn), and *Pseudomonas aeruginosa* (Pae). The dataset included Eco cells exposed to ciprofloxacin for 0, 1, 2, and 4 hours, as well as two species (Pae and Kpn) that were not among the 53 labelled species in the training cohort, providing a test on unlabelled taxa. Clustering of the learned embeddings separated both labelled and unlabelled species and also resolved longitudinal treatment dynamics (Fig. 3c). For Eco, the embeddings distinguished untreated cells (0 hours) from antibiotic-stressed cells (1, 2, and 4 hours), indicating that LucaCell can separate microbial physiological states associated with drug exposure.

Furthermore, the subclusters identified by LucaCell’s alignment-free embeddings for Pae were concordant with those derived from traditional alignment-based gene expression matrices (Fig. 3d, e), as quantified by an Adjusted Rand Index (ARI) of 0.864 and a normalized mutual information (NMI) of 0.785, suggesting that raw-read latent representations can recover biologically meaningful structure comparable to gene-expression-based analyses. Moreover, LucaCell’s alignment-free embeddings achieved higher Silhouette and Calinski-Harabasz scores, and a lower Davies-Bouldin score than gene expression matrices, indicating improved cluster compactness and separation relative to alignment-based expression matrices. Differential expression analysis showed that Pae cluster 0 was enriched for stress-response genes, including *katA*, *azu*, and *arcD*, whereas cluster 1 was dominated by translational machinery and denitrification genes (Fig. 3f). This pattern suggests the coexistence of physiologically distinct subpopulations within a single culture. Taken together, these results indicate that LucaCell provides an alignment-free framework that can both distinguish microbial species and resolve transcriptionally distinct physiological states within species.

### Modeling gene perturbations and variant-associated transcriptomic changes

In-silico simulation of genetic perturbations offers a scalable approach for prioritizing and exploring genetic interactions that would be costly to assess experimentally. To evaluate LucaCell in this domain, we established a sequence-aware predictive framework that integrates cell embeddings extracted from the frozen foundation model with perturbed gene sequence embeddings (Fig. 4a). We benchmarked LucaCell against three state-of-the-art foundation models: Geneformer, scGPT, and scFoundation, using the Adamson CRISPR screening dataset ^22^. LucaCell achieved the lowest Mean Squared Error (MSE) on both differentially expressed genes and global genes, indicating improved reconstruction accuracy for absolute expression values (Fig. 4b). While scGPT achieved the highest mean delta Pearson correlation (0.913) in capturing directional transcriptional trends, LucaCell ranked second (0.851) and outperformed the other baselines (Fig. 4b). These results indicate that LucaCell not only predicts perturbation-induced expression changes with accurate directionality, but also outperforms all benchmarked models in single-cell-level reconstruction, highlighting its superior ability to capture fine-grained expression dynamics.

**Figure 4.**
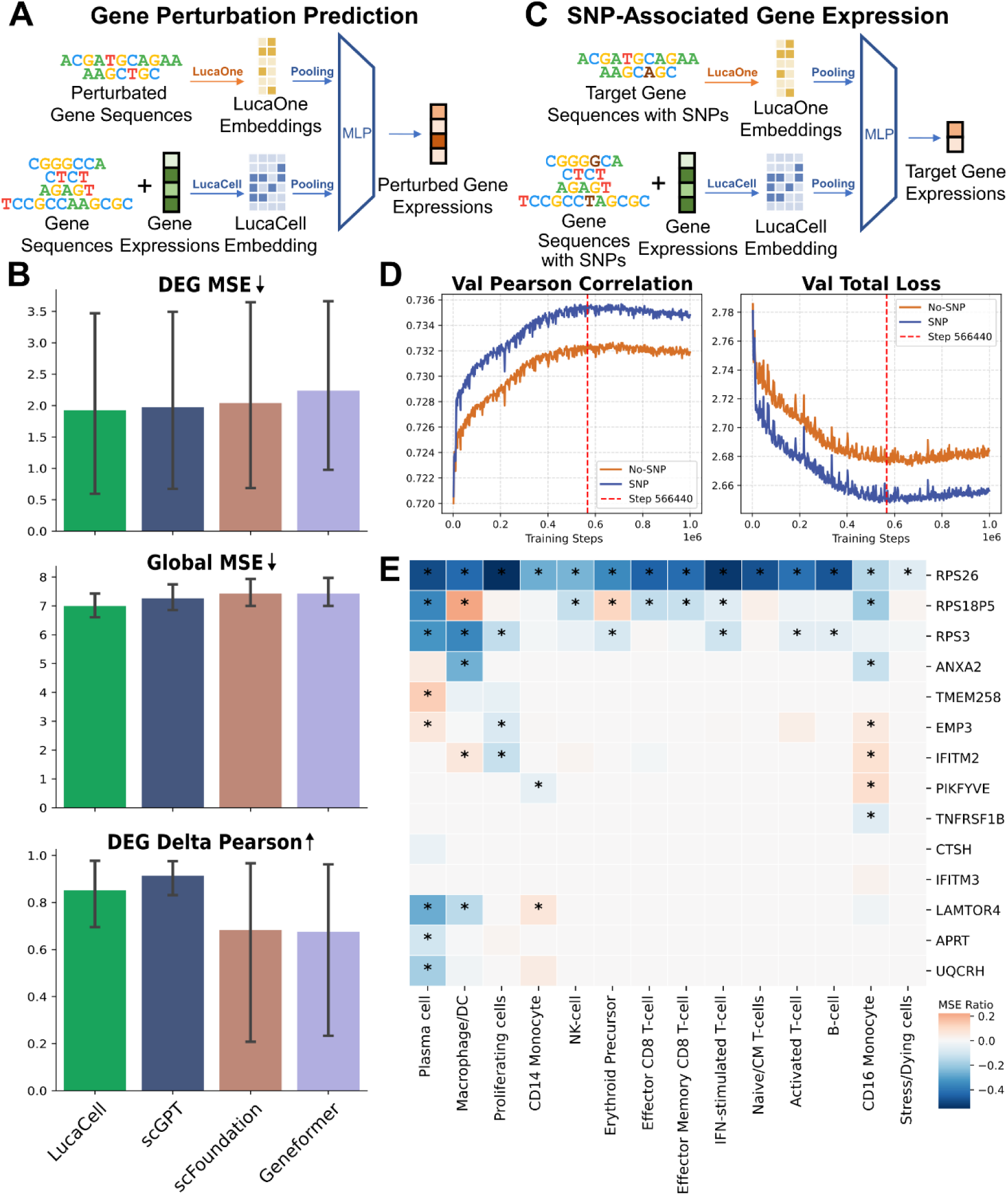
Sequence-aware modeling of gene perturbations and SNP-associated transcriptional changes. a, Schematic of the gene perturbation prediction workflow. The model integrates perturbed gene sequences with baseline cell representation to predict post-perturbation states via an MLP. b, Benchmarking against single-cell foundation model baselines. Comparison of LucaCell, scGPT, scFoundation, and Geneformer across three evaluation metrics: MSE of DEGs (top), global MSE (middle), and Pearson correlation of mean DEG expression changes (bottom). c, Schematic of the SNP-associated gene expression prediction workflow. Target gene sequences harboring specific SNPs are combined with baseline cell representation to predict the expression levels of target genes via an MLP. d, Validation performance and training convergence. Pearson correlation (left) and validation loss (right) profiles for models trained with (blue) and without (orange) donor-specific SNPs. The red dashed line denotes the selected checkpoint at step 566,440. e, Cell type-specific MSE changes upon integrating donor-specific SNPs. Heatmap showing relative changes in MSE across cell types. Only highly expressed genes (detected in ≥ 75% of cells per cell type) were analyzed to mitigate drop-out artifacts. Asterisks (*) denote MSE changes exceeding 5% after integrating SNPs.

While gene-level perturbation modeling provides insights into gene regulatory networks, many disease-associated genetic variations identified by genome-wide association studies (GWAS) are single-nucleotide polymorphisms (SNPs) rather than gene-level loss-of-function events ^23^. To model the potential impact of genomic variation on individual-specific expression, we leveraged the OneK1K dataset ^24^ to predict target gene expression in a cohort of genotyped donors (Fig. 4c). To enable the model to capture nucleotide-level sequence differences, we implemented class token (CLS)-pooled gene-sequence embeddings, and compared the predictive accuracy of donor-specific mRNA sequences harboring exonic SNPs with that of reference sequences. We found incorporating individual-specific SNPs improved global prediction performance (Fig. 4d). This suggests that sequence variation contributes to predictive performance in this setting.

To assess gene-level sensitivity to sequence variation, we compared predictive performance across diverse cell types before and after incorporating SNP information. Incorporating SNPs reduced prediction error (MSE) for ribosomal protein genes, including *RPS26* and *RPS3*, across multiple cell types (Fig. 4e). We also observed lineage-specific patterns, where SNP information selectively improved predictions for genes such as *LAMTOR4*, *APRT*, and *UQCRH* in plasma cells, as well as *ANXA2* in macrophages and CD16 monocytes (Fig. 4e). These observations are consistent with context-dependent effects of SNP on gene expression. Although occasional increases in MSE highlight the difficulty of modeling complex genetic regulatory effects with limited variant-specific data, these findings indicate that LucaCell can leverage sequence variation to improve expression modeling in a gene- and cell type-dependent manner.

### Single-cell viral load prediction

Understanding viral host specificity is essential for elucidating infection mechanisms, predicting spillover risk, and informing intervention strategies. Although predicting viral host ranges at the organismal level is well-established ^25,26^, inferring the infection status of individual cells remains challenging. Single-cell sequencing is inherently destructive, precluding direct longitudinal tracking of viral entry into a given cell; we therefore turned to LucaCell to infer infection status indirectly by examining the transcriptional signatures associated with infection (Fig. 5a). Using virus-infection scRNA-seq datasets ^27–30^ spanning five influenza A virus (IAV) strains (H1N1: PR8, Cal07, Flu-WSN, and Flu-WSN-syn; H3N2: Perth09) across two host cell lines (A549 and LCLs), we predicted single-cell viral load, defined as the ratio of viral read counts to total read counts. To evaluate cross-context generalization, we used a leave-one-combination-out validation scheme, sequentially holding out the Cal07-A549 and Perth09-A549 cohorts as unseen test sets. LucaCell achieved stable viral load prediction, with lower MSE and consistently strong R-squared values and Pearson correlations, compared with baseline models, which exhibited greater variability across viral-host contexts (Fig. 5b). Notably, on the more challenging Cal07 hold-out set, LucaCell was the only model to achieve a positive R-squared, meaning performance superior to the simple mean baseline, whereas all identifier-based baselines (including Geneformer and scFoundation) suffered catastrophic performance collapse (R-squared as low as −1.5), demonstrating that LucaCell’s sequence-based representation of viral genomes confers superior robustness against generalization failure when predicting across genetically divergent, held-out viral strains.

**Figure 5.**
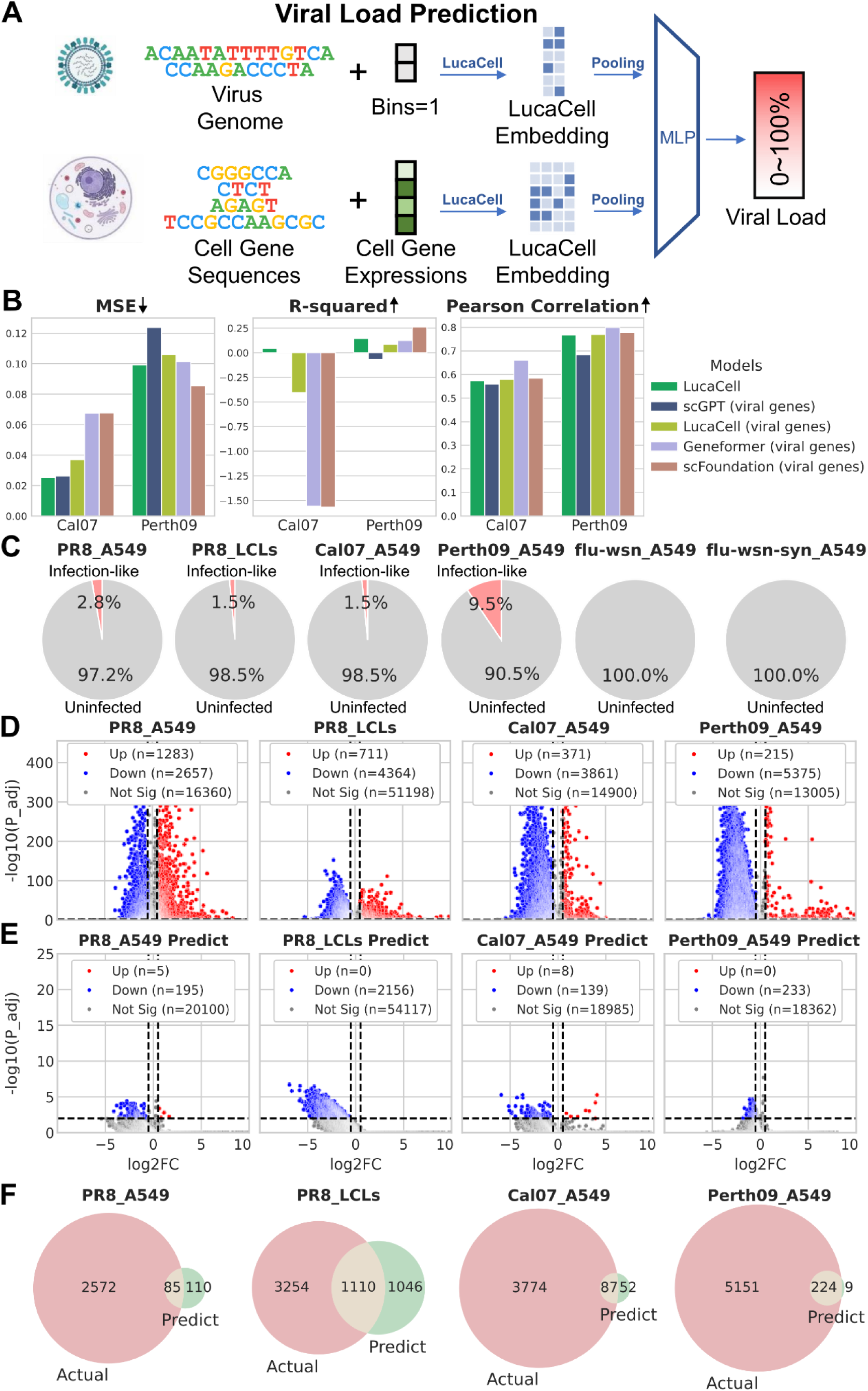
In silico modeling of host-virus interactions at single-cell level. a, Schematic of the sequence-informed viral load prediction framework. The model tokenizes raw viral genomes and host single-cell expression profiles to generate embeddings, which are integrated through an MLP to predict continuous single-cell viral load (0 to 100%). b, Benchmarking against foundation models. Evaluation across MSE (left), R-squared (middle), and Pearson correlation (right) on Cal07 and Perth09 influenza test sets. “LucaCell” (green) represents the joint framework where both host cells and viral genomes are encoded using LucaCell embeddings. The suffix “(viral genes)” denotes baseline configurations where the virus is represented by simply stacking LucaOne gene-level sequence embeddings. LucaCell outperformed LucaCell (viral genes), indicating that viral genomes encoded with LucaCell embeddings are more predictive than those encoded with LucaOne sequence embeddings. c, Proportion of mock-infected cells classified in silico as infection-like (viral load >10%, pink) versus uninfected (viral load ≤10%, grey) across six virus-cell line combinations. d, Volcano plots showing DEGs between experimentally infected (viral load >10%) and uninfected (viral load ≤10%) cells. Red and blue dots represent significant up- and down-regulated genes, respectively (adjusted P < 0.01, |log2FC| > 0.5). e, Volcano plots showing DEGs within the mock-infected pool, comparing cells predicted as infection-like versus uninfected (adjusted P < 0.01, |log2FC| > 0.5). f, Concordance of transcriptional repression between actual infection and predicted infection-like host states. Venn diagrams illustrating the overlap of down-regulated DEGs between experimentally infected (“Actual”, red) and predicted infection-like (“Predict”, green) cells across four virus-cell line combinations.

We next asked whether unexposed control cells contain cells whose baseline transcriptional profile resembles that of infected cells. We applied the model, trained on exposed cells, to unexposed control (mock-infected) cells and scored them with the same regression head. To reduce potential confounding effects, we controlled for mitochondrial gene expression ratio, the number of detected genes, and cell cycle states. Using a 10% predicted viral load threshold to define ‘infected’ cells (Methods), we identified a subpopulation of mock-infected cells displaying infection-like transcriptional signatures (Fig. 5c). Notably, the Perth09-A549 combination showed the highest infection-like cell fraction (9.5%), consistent with previous reports that Perth09-infected cells exhibit significantly higher viral transcript fractions ^29^. These findings suggest that LucaCell is sensitive to strain-specific viral signatures and can identify mock-infected cells with infection-like transcriptional states.

To check this predictive capability, we compared the differentially expressed genes (DEGs) in experimentally infected cells with those in predicted infection-like subpopulation. Both groups showed pronounced and coordinated global downregulation of gene expression (Fig. 5d, e), with significant overlaps among downregulated DEGs (Fig. 5f). This convergence suggests that pre-existing transcriptional states may provide a basis for predicting subsequent infection outcomes. Pathway enrichment analysis of the shared downregulated genes across viral-host combinations revealed pathways involved in antiviral defense, including the stress response ^31^, deubiquitination ^32^, KEAP1-NFE2L2 ^33^, interleukin-12 (IL12) signaling ^34^, and Notch4 signaling ^35^ (Fig. S4). Enrichment of HIV-associated pathways was also observed. Several factors in these pathways, including TASK-1 and APOBEC3G, are known to inhibit HIV replication ^36^, suggesting that their reduced baseline expression may reflect a common infection-associated transcriptional program. We also observed coordinated downregulation of multiple cell cycle-related pathways, pointing to a low-proliferation state. Because host cell cycle arrest at specific phases is a conserved strategy used by viruses, including HIV ^36^ and influenza virus ^37,38^, to facilitate replication, these observations may reflect an infection-permissive cellular state. Finally, pathway enrichment analysis of genes downregulated only in predicted infection-like cells also identified pathways linked to viral infection, including HIV infection, cell cycle, immune system, interferon ^39,40^, and TGF-β ^41^ signaling (Fig. S5). Together, these findings indicate that LucaCell is able to recapitulate transcriptional programs associated with infection, even in the absence of viral exposure.

## Discussion

In this work, we present LucaCell, a sequence-centric foundation model that replaces discrete gene identifiers with mRNA sequence embeddings for single-cell transcriptomics. By embedding genes in a sequence-derived latent space, LucaCell decouples cellular representation from static gene annotations. This design supports robust generalization in predicting gene perturbations and viral load relative to identifier-based frameworks. It also provides a flexible interface for integrating heterogeneous datasets across species and data types while retaining sensitivity to nucleotide-level variation. However, owing to computational constraints, the current implementation uses mean-pooled sequence embeddings as feature representations for pre-training, which may obscure some fine-grained distinctions. Future work could explore base-level sequence representations and context-specific sequence modeling, which could better preserve localized sequence information, capture context-dependent regulatory activity, and improve downstream single-cell foundation models in cell type resolution, perturbation modeling, and cross-modal integration.

Analysis of microbial single cells demonstrates that this sequence-centric representation also generalizes to prokaryotic systems. Despite the sparsity and noise inherent to smRNA-seq data, LucaCell distinguishes both taxonomic identity and intra-species transcriptomic heterogeneity. This capability is biologically and clinically relevant, as it can help characterize transient cellular states associated with antibiotic tolerance, including bacterial persistence, as well as molecular features linked to virulence. By embedding both eukaryotic and prokaryotic cells within a single framework, LucaCell provide a basis for future studies of host-microbe interactions and for cell-level dissection of how human-associated microbes shape host physiology and health.

Like most existing perturbation-prediction frameworks, LucaCell is constrained by the destructive nature of single-cell sequencing, which precludes longitudinal tracking of the same cell before and after intervention. As a result, current models capture population-average responses rather than individual cell trajectories. Integrating emerging clone-tracing technologies ^42^, where lineage-coupled sister cells can be sequentially profiled to capture baseline and perturbed states, may provide a route for training next-generation predictive models on causal regulatory dynamics.

Although LucaCell currently uses mRNA sequences, its modular architecture could be extended. Future work could incorporate non-coding genomic regions or protein-level features. More broadly, biological macromolecules represent only a portion of cellular physiology, as small chemical molecules also critically modulate intracellular processes, as highlighted by recent work bridging genetic and chemical screens from molecular representation to phenotype modeling ^43^. Incorporating chemical compound embeddings together with abundance binning into the LucaCell framework represents a promising direction. More generally, extending LucaCell toward a broader multi-omic framework that integrates molecular and chemical information may improve its ability to represent cellular state.

### Limitations

Although we benchmarked LucaCell against identifier-based foundation models (Geneformer, scGPT, scFoundation), these models also differ in architecture, scale and pre-training corpus, and we did not train a gene-identifier variant of LucaCell under matched architecture, parameters and pre-training data. The reported differences therefore cannot be attributed specifically to the sequence-based gene representation, which we describe as an enabling rather than a demonstrably superior design choice.

## Methods

### Foundation model architecture, pre-training, and checkpoint selection

LucaCell was implemented as a Transformer encoder for sequence-centric single-cell modeling. The model comprised 40 Transformer layers, 40 attention heads (approximately 3B parameters), an embedding dimension of 2,560, and a feed-forward network size of 10,240. Rotary positional embeddings and a final layer normalization were enabled. Attention dropout was set to 0.05 and activation dropout to 0.0. Gene identity was represented by mean-pooled mRNA sequence embeddings extracted from the final-layer output of the LucaOne foundation model, and gene expression values were discretized into 50 uniform non-zero bins ^9^, where 1 denoted the lowest non-zero expression and 50 denoted the highest non-zero expression; zero-expression genes were represented by a special token, [E_NON]. The input for each cell therefore comprised a gene-ordered sequence, with genes arranged by genomic coordinate and each gene represented by a LucaOne-derived mean-pooled sequence embedding and its associated expression bin. During LucaCell training, LucaOne remained frozen and was not updated jointly with LucaCell.

The foundation model was pre-trained on approximately 85 million single-cell transcriptomes, including 67 million human cells and 18 million mouse cells. For each cell, up to 1,000 expressed genes were sampled; if fewer than 1,000 expressed genes were detected, all expressed genes were retained. Zero-expression genes were then sampled up to the minimum of four times the number of expressed genes, the remaining capacity to a fixed input length of 1,200 tokens, and the total number of available zero-expression genes. This sampling scheme preserved background gene context while constraining the total number of input tokens. Selected genes were ordered by genomic coordinate.

During pre-training, random masking was applied to all sampled genes. The model was trained using a masked gene modeling objective to predict masked expression bins from the surrounding cellular context. Optimization was performed with AdamW (*β*_1_ = 0.9, *β*_2_ = 0.98, weight decay = 0.01) and a peak learning rate of 2 × 10⁻⁴. Training was conducted for up to 3 epochs using distributed data parallelism across 8 Nvidia A100 GPUs, with a per-GPU batch size of 1 and gradient accumulation over 32 steps, yielding an effective batch size of 256. A linear warmup of 32,000 steps was used, and the maximum number of optimization steps was set to 10,000,000. Gradient clipping was applied with a maximum norm of 1.0. Mixed-precision training was performed in bfloat16. A fixed random seed of 1111 was used for reproducibility.

Because pre-training was self-supervised and intended to learn a general-purpose backbone rather than optimize a task-specific objective, we selected the 6.4M-step checkpoint, where the training loss had reached a stable plateau and further improvements were marginal. This checkpoint was used for downstream analyses, feature representation extraction, and fine-tuning initialization.

### Datasets used for downstream analyses

Detailed dataset accessions are listed in Supplementary Table S2.

### Settings for downstream tasks

Task-specific hyperparameters are summarized in Supplementary Table S3.

### Cross-species cell type prediction

The datasets comprised human scRNA-seq and scATAC-seq data from GSE254185 ^14^, as well as lemur ^15^ and mouse ^16^ scRNA-seq data from CELLxGENE ^3^. None of these cohorts were included in pre-training. Genes longer than 10,240 bp were excluded based on hg38, mm10, and micMur2 annotations; for human scATAC-seq, peaks longer than 10,240 bp or mapped to non-standard scaffolds were removed. At the cell level, low-quality cells with fewer than 200 detected genes in scRNA-seq or fewer than 800 accessible peaks in scATAC-seq were filtered out. Cell labels were harmonized across cohorts to standardized consensus cell type annotations.

For cell tokenization, genes and chromatin peaks were ordered by chromosomal coordinate, and the top 2,048 expressed genes for scRNA-seq and top 8,192 accessible peaks for scATAC-seq were selected and discretized into 50 bins. We used a species-held-out split, with human and lemur cells partitioned into training, validation and test sets at an 80%:10%:10% ratio, which were used for model training, final checkpoint selection and performance reporting, respectively, and mouse cells reserved as an additional independent test cohort.

The downstream classification model used a single-layer transformer encoder followed by value-attention pooling ^44^ and a fully connected classification head. The embedding layer was initialized from the pre-trained LucaCell checkpoint and updated during training. Models were optimized with weighted categorical cross-entropy using AdamW, using the hyperparameters as described above.

Cell embeddings were extracted from the pooling layer (the representation vectors) and visualized by UMAP using Scanpy ^45^. We computed centroid embeddings by averaging cell embeddings within each origin group, and calculated pairwise Euclidean distances between origin-level centroids.

For cross-species marker conservation, marker genes were identified independently in human, mouse and lemur for each cell type using Wilcoxon rank-sum tests in Scanpy, restricted to shared orthologous genes, and the top 20 markers per cell type were retained for each species. Marker overlaps were quantified by intersecting species-specific marker sets and visualized with Venn diagrams at both the cell type level and after merging markers across cell types.

Orthologous marker sequence embeddings were generated with LucaOne, and pairwise cosine distances were computed for conserved marker genes across human, mouse and lemur. Distance distributions for Human-Mouse, Human-Lemur and Mouse-Lemur comparisons were evaluated using paired two-sided t-tests.

### Alignment-free single-microbe embedding

We retrieved smRNA-seq datasets PRJCA017256 (human gut) ^20^ for training and CRA011274 (bacterial culture) ^21^ for independent validation. We downloaded the reference genome of Pae (*Pseudomonas aeruginosa*, GCA_000006765) from GenBank. Raw paired-end reads from smRNA-seq datasets were processed with UMI-tools (v1.1.6) ^46^ in extract mode to obtain a 20-bp cell barcode and an 8-bp UMI from read 1, which were appended to the header of read 2. Only read 2 sequences containing cDNA inserts were retained for training and analysis.

Taxonomic profiling was performed with MICtools (v1.0.0) ^47,48^ against the UHGG database (v2.0.1) ^49^, and the proportion of reads assigned to the dominant species within each barcode was used as the taxonomic confidence ratio.

Barcodes with UMI counts between 100 and 2,000 were retained, and reads with Phred quality scores below 20 or ambiguous base fractions above 10% were discarded. We then applied stochastic reservoir sampling to retain up to 8,192 high-quality reads per barcode. Cells with a taxonomic confidence ratio less than 50% (less than 50% of reads were assigned to the dominant species), or belonging to species represented by fewer than 20 cells, were assigned to an ‘unidentifiable’ category. For species represented by more than 100 cells, only the 100 cells with the highest confidence ratios were retained.

To adapt the model to microbial data, sampled reads were used in place of transcripts in random order without genomic alignment, and because read abundance was not used as a biological signal in the alignment-free setting, all reads were assigned to a uniform expression bin of 1, to encode read presence in the absence of expression quantification, reflecting the fact that each RNA-seq read contributes a single count as the minimal unit of expression. The curated dataset comprised 54 taxonomic classes and was split into training, validation and test sets using an 80%:10%:10% ratio, which were used for model training, final checkpoint selection and performance reporting, respectively. Model training followed the protocol used for cross-species cell type prediction, using the hyperparameters as described above. Unsupervised clustering of the 2,560-dimensional cell embeddings was performed in Scanpy using 50 principal components (PCs) and Leiden clustering at resolution 2.0, followed by UMAP visualization.

For Pae expression matrix validation, all qualified reads were aligned to the reference using STAR (v2.7.10a) ^50^, and gene-level counts were generated with featureCounts (v2.0.1) ^51^. Aligned reads were sorted and indexed with Samtools (v1.21) ^52^ to construct the single-microbe expression matrix.

Cluster quality for LucaCell embeddings and gene expression-based representations was assessed using the first 50 PCs of each representation, with the Silhouette score, Calinski-Harabasz index and Davies-Bouldin index. Concordance between clustering results was quantified using ARI and NMI.

### Gene perturbation prediction

We utilized the Adamson dataset (GSE90546) ^22^, and selected the top 2,000 highly variable genes. We developed a sequence-aware perturbation modeling framework for benchmarking single-cell foundation models, in which the perturbed gene was encoded by its LucaOne sequence embedding instead of being included as an input expression feature. To avoid trivial inference from the loss of target-gene expression, all candidate perturbed genes were excluded from cell feature set across all perturbation conditions.

For cell tokenization in LucaCell, raw gene counts were normalized to logCPM (log-transformed counts per million), and highly variable genes with non-zero expression were retained as inputs to match the settings used for the other foundation models. For the baseline models (Geneformer v2 ^7^, scGPT ^9^ and scFoundation ^8^), data were processed using their default tokenization protocols. In all cases, embeddings were extracted from the frozen foundation model.

We used a gene-level out-of-distribution split, with perturbed target genes divided into training, validation and test sets at an 80%:10%:10% ratio, which were used for model training, final checkpoint selection and performance reporting, respectively. Unperturbed control cells were split using the same ratio, and each perturbed cell was randomly paired with a control cell from the corresponding split. The input consisted of the perturbed-gene sequence embedding and the control-cell embedding, and the target was the logCPM expression profile of the perturbed cell.

We trained a pairwise multi-target regression model to predict the expression levels of highly variable genes from paired cell embedding matrix and perturbed target gene embedding vectors. Pairwise regression was performed with a heterogeneous matrix encoder followed by value-attention pooling, addition-based fusion with the gene embeddings, and a fully connected prediction head. Models were optimized with AdamW and a dynamically weighted MSE loss (logCPM threshold, 0.5; weights, 1 and 5 for low- and high-expression targets, respectively).

Performance was evaluated using DEG Delta Pearson, DEG MSE and global MSE. For DEG Delta Pearson, the top 20 DEGs for each perturbation were identified by comparing perturbed and control cells, and Pearson correlation was computed between the predicted and observed average expression changes ^9,10^. DEG MSE and global MSE were calculated for each perturbed cell on DEGs only or across all highly variable genes, respectively.

### Personalized gene expression modeling

We used genotype and single-cell expression profiles from the OneK1K dataset ^24^. Because the genotyping matrices provided per-individual allele dosages rather than phased haplotypes, we constructed allele-frequency-weighted personalized embeddings for background expressed genes and target eGenes (genes regulated by SNPs). Firstly, for each exonic SNP site, allele-specific sequences were generated by substituting each allele into the reference gene sequence, and LucaOne CLS embeddings were extracted for each allele-specific sequence. Then individual-level personalized embeddings were approximated by weighting allele-specific embeddings by the observed per-individual allele dosages and aggregating across exonic SNPs. Exonic eSNPs (SNPs that regulate gene expressions) and their target eGenes were obtained from the OneK1K eQTL map ^24^. We retained eGenes linked to exonic eSNPs, and used their expression levels (logCPM) as regression targets.

To reduce sex chromosome-related confounding, we restricted the analysis to autosomal genes. To prevent information leakage, target eGenes were excluded from the cell token set, and cells were tokenized using the top 2,048 background expressed genes under two conditions: a reference setting in which genes were mapped to wild-type LucaOne embeddings, and a SNP-aware setting in which genes were mapped to personalized SNP-weighted embeddings. The two settings differed only in the source of gene embeddings; in both cases, tokenized cell profiles were processed by the frozen LucaCell foundation model to obtain cell embeddings.

We split unique donors into training, validation and test sets at an 80%:10%:10% ratio, which were used for model training, final checkpoint selection and performance reporting, respectively, and trained a pairwise multi-target regression model to predict the expression levels of multiple eGenes from paired cell and eGene embeddings. Pairwise training matched each cell embedding matrix with the corresponding set of eGene sequence embeddings under two matched conditions: in the SNP-aware setting, both cell and eGene representations were based on personalized SNP-weighted embeddings, whereas in the reference setting, both were based on wild-type embeddings. Model training followed the protocol used for gene perturbation prediction, using the hyperparameters as described above.

To evaluate the effect of personalized exonic SNPs, predicted eGene expression values from the SNP-aware and reference models were compared with the ground truth in the test cohort. Analysis was restricted to genes expressed in at least 75% of cells within each cell type. Performance was quantified using MSE, and the relative change in MSE was calculated as 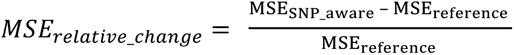.

### Viral load prediction

We compiled single-cell RNA-seq datasets from four cohorts spanning multiple influenza A virus (IAV) strains and host cell lines: GSE122031 (PR8 in A549 cells) ^28^, GSE143167 (Cal07 and Perth09 in A549 cells) ^29^, GSE205796 (PR8 in lymphoblastoid cell lines) ^30^, and the Russell cohort (wild-type flu-wsn and flu-wsn-syn in A549 cells) ^27^. Genomic references for PR8 (NCBI Taxonomy ID 211044), Cal07 (NCBI Taxonomy ID 641809) and Perth09 (NCBI Taxonomy ID 654811) were obtained from the NCBI Virus database, and reference sequences for flu-wsn and flu-wsn-syn were obtained from the Russell repository ^27^.

Viral load was defined as the ratio of the summed counts of the eight core viral genes (*PB2*, *PB1*, *PA*, *HA*, *NP*, *NA*, *M* and *NS*) to the total cellular transcript count for each cell 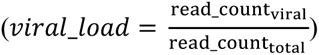. Cells with fewer than 100 detected host genes were excluded.

Cells were tokenized using the top 2,048 expressed host genes, which were discretized into 50 bins. The frozen LucaCell foundation model was used to generate cell embeddings, whereas baseline models (Geneformer v2, scGPT and scFoundation) were processed using their native tokenization protocols.

To represent the viral genome, we used two schemes. For baseline models, the eight viral genes were ordered as (*PB2*, *PB1*, *PA*, *HA*, *NP*, *NA*, *M* and *NS*), and their LucaOne sequence embedding vectors were stacked into a matrix. For LucaCell, the eight viral genes were represented as tokens in the same order, with all genes assigned an expression bin of 1, to represent viral presence in the absence of endogenous expression quantification.

To assess generalization to unseen viral lineages, we trained two strain-level hold-out models in which A549 cells infected with Cal07 or Perth09 were reserved as independent test sets for performance reporting, respectively. The remaining cells were split into training and validation sets at an 80%:10% ratio, while maintaining a balanced distribution of virus-cell line combinations. We also trained a unified multi-strain model on all viral strains and both cell lines to predict infection-like states on corresponding mock control cells.

We trained a pairwise regression model to predict viral load from paired cell and viral genome embedding matrices. Pairwise models used a shared architecture with linear input projection, a dual-branch transformer matrix encoder, value-attention pooling and a regression head. Models were optimized with AdamW, using the hyperparameters as described above. Performance was evaluated using Pearson correlation, R-squared and MSE.

We evaluated four cutoffs (1%, 5%, 10% and 20%) for classifying cells as infected. The 1% and 5% cutoffs yielded a higher proportion of predicted infected cells in mock samples than in experimentally infected samples for some virus-cell line combinations, whereas the 20% cutoff failed to detect infected cells in some infected datasets. We therefore selected the 10% cutoff, which balanced sensitivity and specificity.

To validate infection-like predictions in mock cells, we performed differential expression analysis using Wilcoxon rank-sum tests in Scanpy. Mock cells with mitochondrial read fractions above 10% or total detected genes less than two standard deviations below the dataset mean were excluded. We calculated G1, S and G2/M phase scores and assessed cell-cycle phase distributions using chi-square tests. No significant differences (P > 0.05) in the proportions of G1, S and G2/M cells were observed between predicted infection-like and uninfected mock cells across the PR8_A549, PR8_LCLs, Cal07_A549, and Perth09_A549 combinations.

We identified DEGs separately in experimentally infected cells and predicted infection-like mock cells using the criteria of adjusted P < 0.01 and absolute log2 fold change > 0.5. Pathway enrichment analysis were performed using GSEApy ^53^ against the Reactome 2022 database ^54^.

## Data availability

The pre-training dataset of LucaCell is available at http://47.93.21.181/lucacell/. This IP-based link is provided as a temporary access point because the dataset is too large for currently available public repositories; we will attempt to deposit it in an appropriate public database in the future. The datasets of all downstream tasks and other supplementary materials are available at https://zenodo.org/records/22073984.

## Code availability

The source code for LucaCell pre-training, embedding inference, and downstream tasks is available at https://github.com/LucaOne/LucaCell. The trained model weights are available at https://huggingface.co/LucaGroup/LucaCell-v1.0-step6.4M.

## Acknowledgements

We thank Q. Kong and X. Xia for insightful discussions and comments on the manuscript. Y. Sun gratefully acknowledges BGI for support during their training. We are grateful to Alibaba Cloud for providing computational resources and support during model training and analysis. We also acknowledge publicly available single-cell data resources, such as CZ CELLxGENE, GEO, and OneK1K, which made this study possible.

## Author contributions

Y. Sun., Y. He, H. Yang, and Z. Wang conceived the study. Y. Sun and Y. He designed the LucaCell framework. Y. He developed and implemented the model. Y. Sun and M. Ren constructed the training dataset and performed data preprocessing. Y. He carried out model pretraining and benchmarking. Y. Sun, Y. He, Y. Wang, and P. Xu performed downstream biological analyses and interpreted the results. Y. Sun and Y. He drafted the manuscript. Y. Hou, Y. Kang, T. Hou., and J. Ye revised the manuscript. Y. He, H. Yang, and Z. Wang supervised the study. All authors contributed to manuscript revision and approved the final version.

