## Supplementary Information for "LucaCell: a sequence-centric foundation model for cross-species single-cell analysis"

### Supplementary Materials

### Supplementary Figures

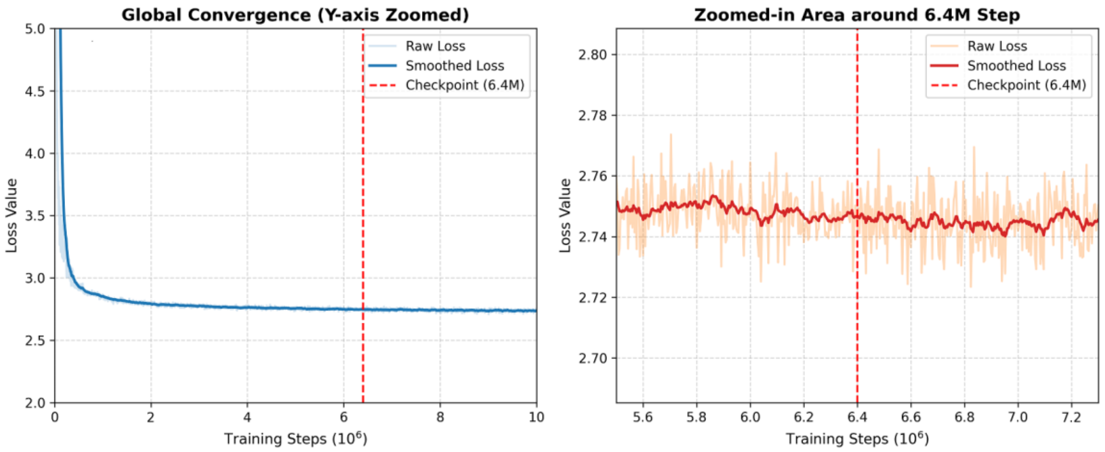


Figure S1. Pre-training loss trajectory of LucaCell. Global self-supervised pre-training loss trajectory across 10 million steps (left), with a zoomed-in view around the 6.4 million (M) step mark (right). The red dashed line indicates the 6.4M-step checkpoint selected for downstream tasks and fine-tuning initialization.

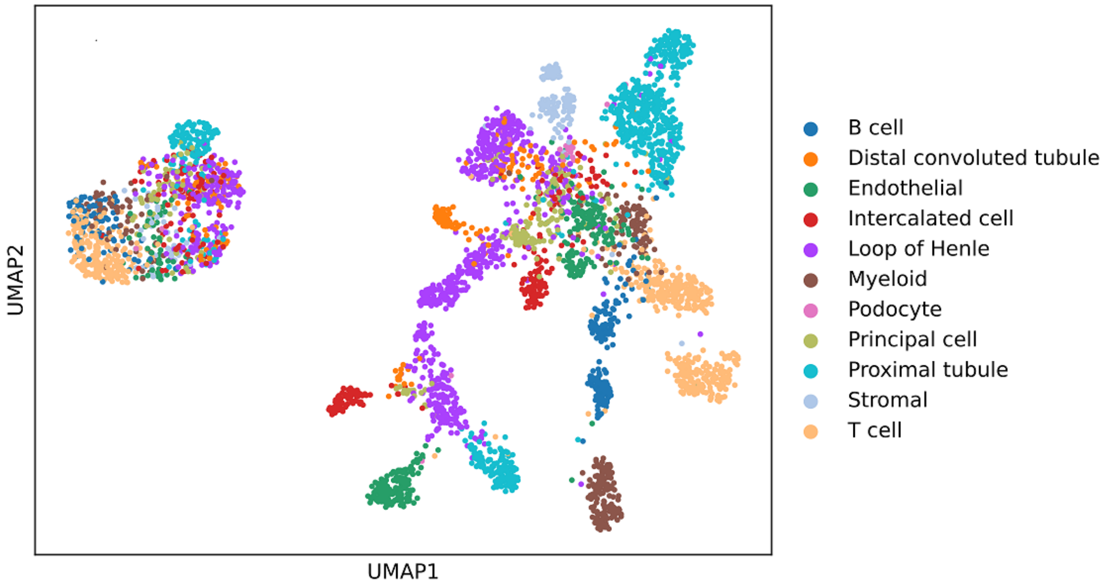


Figure S2. UMAP visualization of the joint latent space containing cross-species and cross-modal cell representations, colored by cell types.

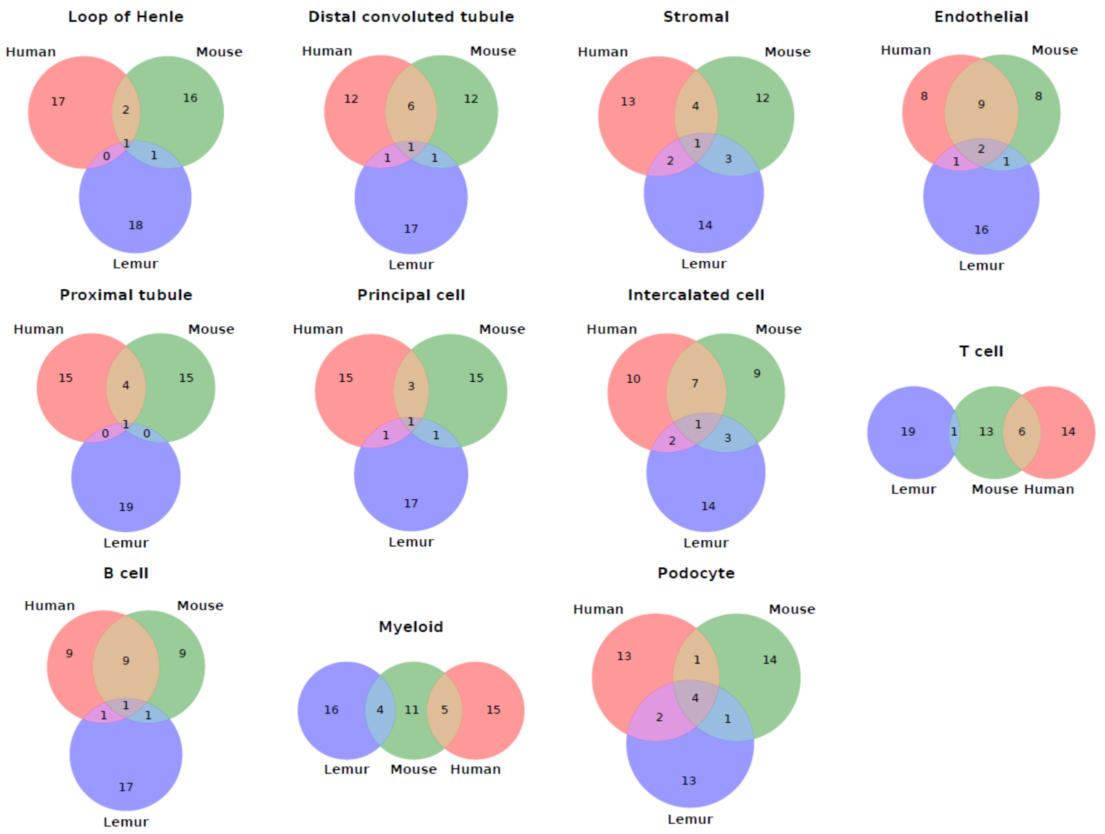


Figure S3. Overlap of marker genes among human, mouse, and lemur, shown separately for each cell type.

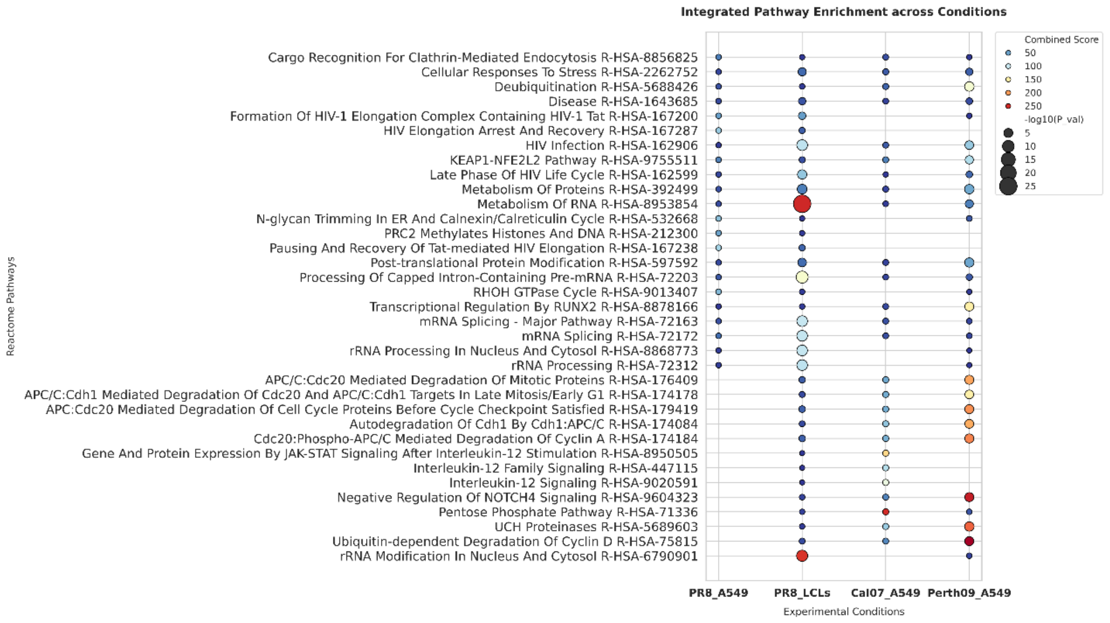


Figure S4. The enriched Reactome pathways for the overlapping down-regulated DEGs across four virus-cell line combinations. For each combination, the top 10 enriched pathways were selected and merged for visualization.

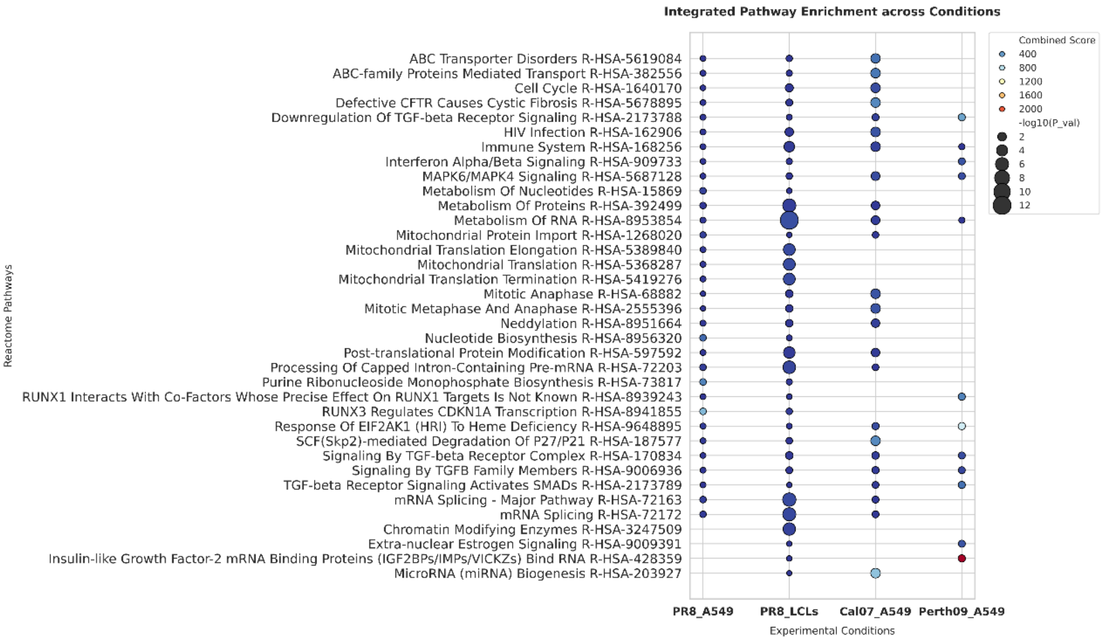


Figure S5. The enriched Reactome pathways for the down-regulated DEGs genes detected exclusively in predicted infection-like cells. The top 10 enriched pathways were selected and merged for visualization.

**Supplementary Tables**

Table S1. Comparison among single-cell foundation models

| Model | Gene representation | Expression representation | Cross-species transfer | Non-coding region inclusion | Nucleotide-level mutation sensitivity | RNA-seq read-level input |
|---|---|---|---|---|---|---|
| Geneformer [1] | Gene ID/Name | Ranked expression | ✗ | ✗ | ✗ | ✗ |
| scFoundation [2] | Gene ID/Name | Continuous expression | ✗ | ✗ | ✗ | ✗ |
| scGPT [3] | Gene ID/Name | Binned expression | ✗ | ✗ | ✗ | ✗ |
| CellFM [4] | Gene ID/Name | Continuous expression | ✗ | ✗ | ✗ | ✗ |
| TranscriptFormer [5] | Protein sequence-derived embedding | Raw counts | ✓ | ✗ | ✗ | ✗ |
| UCE [6] | Protein sequence-derived embedding | Count-weighted sampling | ✓ | ✗ | ✗ | ✗ |
| LucaCell | mRNA sequence-derived embedding | Binned expression | ✓ | ✓ | ✓ | ✓ |

Table S2. Datasets used for downstream analyses

| Task | Dataset | Source | Accession |
| --- | --- | --- | --- |
| Cross-species cell type prediction | Human kidney scRNA-seq and scATAC-seq [7] | GEO [8] | GSE254185 |
| | Lemur kidney scRNA-seq [9] | CELLxGENE [10] | cellxgene.cziscience.com/collections/a137437b-d284-4a27-b1e9-36958a8f92c1 |
| | Mouse kidney scRNA-seq [11] | CELLxGENE | cellxgene.cziscience.com/collections/9c9d04c4-8899-417f-bb6f-6107dcadf14f |
| Alignment-free microbe embedding | Human gut smRNA-seq [12] | Genome Sequence Archive [13] | PRJCA017256 |
| | Bacterial culture smRNA-seq [14] | Genome Sequence Archive | CRA011274 |
| Gene perturbation prediction | Adamson [15] | GEO | GSE90546 |
| SNP-associated gene expressions | OneK1K [16] | GEO | GSE196830 |
| Viral load prediction | PR8_A549 [17] | GEO | GSE122031 |
| | Cal07_ A549 and Perth09_ A549 [18] | GEO | GSE143167 |
| | PR8_LCLs [19] | GEO | GSE205796 |
| | Russell [20] | / | datadryad.org/dataset/doi:10.5061/dryad.qp0t3 |

None of the datasets used for downstream tasks were included in the pre-training corpus.

Table S3. Task-specific hyperparameters for downstream tasks

| Tasks | Maximum epochs | Peak learning rate | Batch size | Warmup steps | Loss Type | Best metric |
|---|---|---|---|---|---|---|
| Cross-species cell type prediction | 80 | $2\times10^{-4}$ | 16 | 1,000 | Cross entropy | Macro-F1 |
| Alignment-free microbe embedding | 100 | $2\times10^{-4}$ | 16 | 1,000 | Cross entropy | Macro-F1 |
| Gene perturbation prediction | 200 | $1\times10^{-4}$ | 16 | 1,000 | Global MSE | Pearson correlation |
| SNP-associated gene expressions | 300 | $1\times10^{-4}$ | 16 | 1,000 | Global MSE | Pearson correlation |
| Viral load prediction | 500 | $1\times10^{-5}$ | 16 | 1,000 | Global MSE | Pearson correlation |

'Loss Type' denotes the training objective used during model building, and 'Best metric' denotes the validation metric used to select the best checkpoint at the epoch level for performance reporting and inference. For all comparative experiments, the models were trained and evaluated using identical hyperparameter settings and random seeds, and all downstream tasks were conducted on a single NVIDIA A100 GPU. Unless otherwise noted, all downstream tasks used the hyperparameters summarized here.
